# Modeling human retinal ganglion cell axonal outgrowth, development, and pathology using pluripotent stem cell-based microfluidic platforms

**DOI:** 10.1101/2025.05.02.651934

**Authors:** Cátia Gomes, Kang-Chieh Huang, Sailee S. Lavekar, Jade Harkin, Carson Prosser, Yue Fang, Claire Kalem, Adrian Oblak, Chi Zhang, Jason S. Meyer

## Abstract

Retinal ganglion cells (RGCs) are highly compartmentalized cells, with long axons serving as the sole connection between the eye and the brain. RGC degeneration in injury and/or disease also occurs in a compartmentalized manner, with distinct injury responses in axonal and somatodendritic compartments. Thus, the goal of this study was to establish a novel microfluidic-based platform for the analysis of RGC compartmentalization in health and disease states. Human pluripotent stem cell (hPSC)-derived RGCs were seeded into microfluidics, enabling the recruitment and isolation of axons apart from the somatodendritic compartment. Initial studies explored axonal outgrowth and compartmentalization of axons and dendrites. We then compared the differential response of RGCs differentiated from hPSCs carrying the OPTN(E50K) glaucoma mutation with isogenic control RGCs in their respective axonal and somatodendritic compartments, followed by analysis of axonal transport. Further, we explored the axonal transcriptome via RNA-seq, focusing on disease-related axonal differences. Finally, we established models to uniquely orient astrocytes along the axonal compartment combined with modulation of astrocyte reactivity as a pathological feature of neurodegeneration. Overall, RGC culture within microfluidic chips allowed enhanced cell growth and maturation, including long-distance axonal projections and proper compartmentalization, while patient-specific RGCs exhibited axonal outgrowth deficits as well as decreased rate of axonal transport. Finally, the induction of astrocyte reactivity uniquely along the proximal region of RGC axons led to the onset of neurodegenerative phenotypes in RGCs. These results represent the first study to effectively recapitulate the highly compartmentalized properties of hPSC-derived RGCs in healthy and disease states, providing a more physiologically relevant in vitro model for neuronal development and degeneration.

## INTRODUCTION

Retinal ganglion cells (RGCs) are the projection neurons of the retina, serving as the sole connection between the eye and the brain (1, 2). RGCs extend long distance axonal projections along the optic nerve from the retina, mostly to thalamic targets in the brain. Owing to these long-distance projections, RGCs are also highly compartmentalized cells, with several unique features specific to RGC axons compared to other RGC regions (3, 4). Additionally, the axonal region of RGCs is of considerable relevance to many diseases and injuries of the optic nerve including glaucoma and traumatic brain injury, as the axon is the primary site of injury in these conditions (5). As such, several studies have focused upon the investigation of cellular mechanisms underlying RGC axonal function in healthy and disease states, as well as strategies to promote regeneration and neuroprotection of RGC axons in the optic nerve (6–8). However, despite the importance of RGCs and their axons extending through the optic nerve, it has been relatively difficult to properly investigate human RGC axons, given their relative inaccessibility to experimental modulation and limited regenerative capabilities.

Human pluripotent stem cells (hPSCs) offer a powerful alternative to the investigation of human RGCs, as they can be derived from specific patients or engineered to harbor disease-associated genetic variants, have the potential for virtually unlimited expansion, and can be directed to differentiate into any cell type, including RGCs. Indeed, many studies over the past decade have focused upon the derivation of RGCs from hPSCs (9–14), including applications for studies of human retinal development as well as the *in vitro* modeling of RGC neurodegenerative diseases such as glaucoma (10, 11, 14). Often, these studies have focused on RGCs as a whole, without regard for the highly compartmentalized nature of RGCs nor the importance of RGC axons to the neurodegenerative state associated with glaucoma or other RGC-associated neurodegenerative conditions. As such, a critical need exists to develop advanced models and experimental approaches with which to better analyze RGC axons *in vitro*, which would result in a greater understanding of some of the mechanisms by which human RGC axons degenerate in disease states, as well as offering an opportunity to investigate strategies that promote human RGC axonal outgrowth and regeneration.

To address these shortcomings, in this study we sought to advance the investigation of RGC axons through the implementation of microfluidic platforms for the growth of RGCs and recruitment of RGC axonal outgrowth. hPSC-derived RGCs were seeded into microfluidic platforms, with experimental chambers connected through microgrooves that permit axonal outgrowth yet prevent migration of cell bodies. As such, RGCs could be seeded into one chamber of the microfluidic platform, and their axons could be effectively isolated within the other corresponding microfluidic chamber. Through these studies, we found that RGCs became much more polarized and exhibited greater compartmentalized features compared to the same cells grown in traditional 2D cultures. Given the ability to specifically investigate RGC axons, we then assessed transcriptional differences within axonal compartments through Axon-Seq approaches, revealing numerous axonally-regulated transcripts. As RGC axons are the initial site of injury in diseases such as glaucoma, we then sought to identify how axons are modulated in conditions relevant to glaucoma by using isogenic cell lines with the glaucoma-associated OPTN(E50K) mutation as well as a corresponding control cell line, demonstrating deficits specifically within OPTN(E50K) axons including total outgrowth as well as axonal transport. Finally, as the proximal region of RGC axons are a specific site of injury in glaucoma, we then advanced these microfluidic approaches to mimic neuroinflammatory responses related to those found within the optic nerve head by co-culture of hPSC-derived astrocytes along the proximal RGC axonal region using three-chamber microfluidic platforms, demonstrating that glial activation specifically along the proximal axonal region resulted in neurodegenerative phenotypes along RGCs. Taken together, the results of this study underscore the importance for specifically investigating RGC axons in disease states and provide the powerful foundation and justification for future studies targeting RGC axons for disease modeling.

## RESULTS

### Microfluidic devices promote hPSC-RGCs polarization with somatodendritic and axonal specification

hPSC-derived RGCs are powerful in vitro models to investigate human RGCs, either for studies of human retinal development or disease-associated RGC degeneration. However, prior studies have mainly been performed in a simple culture system that fails to account for the highly compartmentalized nature of RGCs, including defined somatodendritic and axonal compartments. (15, 16). Microfluidic platforms have been shown to be capable of isolating axonal compartments of neuronal cells (17, 18), providing a powerful approach for studies of how the RGC axonal compartment is specifically modulated during developmental maturation as well as during injury or disease. To better understand how RGC neurites develop in microfluidic devices, we compared the axonal outgrowth of hPSC-RGCs in standard cultures or microfluidic platforms, respectively (Figure 1). Initially, hPSC-RGCs were maintained in standard cultures for 4 weeks to allow for maturation, and neurites were then stained with antibodies against MAP2 and SMI-312, markers of the somatodendritic and axonal compartments, respectively. hPSC-RGCs grown in standard culture conditions exhibited neurite elongation and complexity, indicating RGC morphological maturation similar to our previous studies (Figure 1B-E)(11, 19–21). However, hPSC-RGCs grown in standard cultures demonstrated only limited dendritic and axonal polarization, as some neurites were positive for both MAP2 and SMI-312 (white arrowheads), while other neurites solely expressed either MAP2 or SMI-312 (red or cyan arrowheads, respectively) (Figure 1C-E). Interestingly, when hPSC-RGCs were seeded into microfluidic devices, we observed a more robust separation of MAP2 and SMI-312, suggesting greater refinement of somatodendritic and axonal compartments (Figure 1G-J). hPSC-RGC neurites in the somatic area seemed to exhibit only marginal co-localization between MAP2 and SMI-312, with MAP2 positive neurites restricted to the somatic chamber and surrounding the RGC cell body, morphologically more typical of dendrites (Figure 1H-J). On the other hand, hPSC-RGC neurites in the axonal chamber exhibited a robust and exclusive SMI-312 expression, with undetectable MAP2 staining (Figure 1G), confirming that the microgrooves connecting the two chambers in the microfluidic platforms only allowed axons to cross through. Focusing on the somatodendritic compartment of RGCs (Figure 1K-T), we observed a significant increase in the number of primary MAP2-positive dendrites in RGCs grown on microfluidic devices compared to standard cultures (Figure 1S), further demonstrating that RGC polarization in microfluidic systems also contributes to a faster maturation and complexity. Moreover, Imaris imaging software was used to more precisely quantify the extent of co-localization of MAP2 and SMI-312 (Figure 1K-R), in which we observed greatly decreased co-colocalization of MAP2 and SMI-312 in the somatic region of RGCs grown on microfluidic devices compared to standard cultures (Figure 1T), providing further evidence of greater RGC maturation when grown in microfluidic devices. Collectively, these results confirmed the greater utility of microfluidic devices as a stronger experimental approach to induce RGC maturation and polarization in a more physiologically relevant manner.

**Figure 1.**
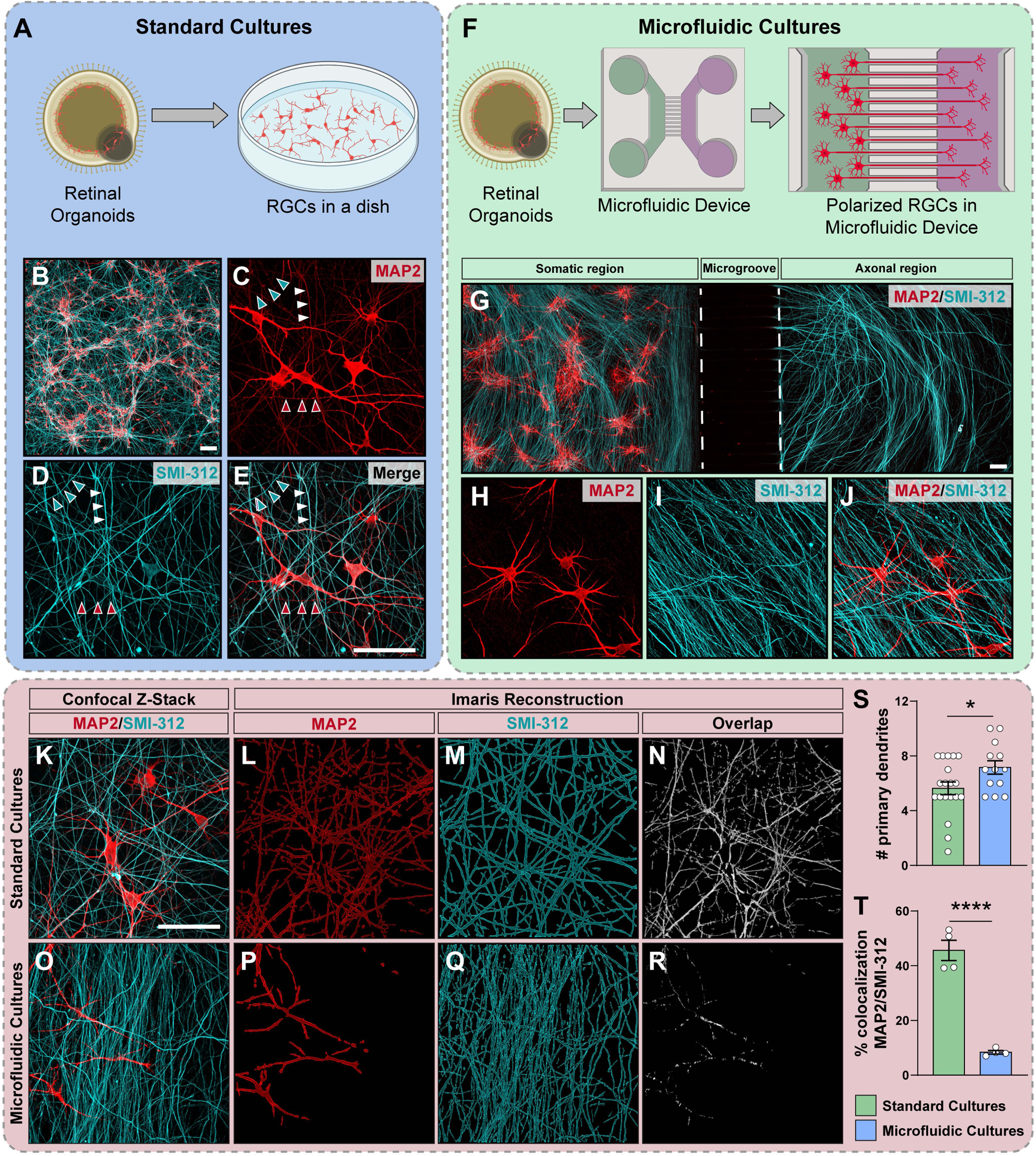
hPSC-derived RGCs demonstrate somatodendritic and axonal specification when seeded in microfluidic devices. (A) Schematic of standard cultures of hPSC-derived RGCs seeded on a dish. (B) Representative images of hPSC-RGC neurite outgrowth seeded in coverslips. (C-E) High magnification images show MAP2-positive dendrites (red arrowhead) and SMI-312 positive axons (green arrowhead) in hPSC-RGCs, while some neurites (white arrowhead) exhibit incomplete specification with co-staining of both MAP2 and SMI-312. (F) Schematic representation of RGCs seeded in microfluidic platforms. (G) Representative images of hPSC-RGC neurite outgrowth in microfluidic chips. (H-J) High magnification images show a nearly complete specification in somatic and axonal chamber with scarce colocalization between MAP2 and SMI-312 in neurites. (K-N) Representative images of standard cultures of RGCs and respective Imaris model for MAP2 and SMI-312 staining as well as co-localization of both. (O-R) Representative images of RGCs seeded in the somatic chamber of microfluidic platforms and respective Imaris model for MAP2 and SMI-312 staining as well as co-localization of both. (S) Number of primary MAP2-positive dendrites in standard and microfluidic cultures. (T) Percent of MAP2 staining co-localized with SMI-312 in both standard and microfluidic cultures, determined by Imaris. Data represent mean values ± S.E.M. *pL<L0.05, ****pL<L0.0001 *vs.* standard cultures, two-tailed unpaired Student’s t test. Scale bar: 100 μm across all images.

Given the ability to effectively isolate RGC axons into the axonal compartment of microfluidic devices in the absence of any RGC somas or dendrites, we then sought to explore the localization of RNA transcripts selectively within developing axons (Figure 2). As long-distance projection neurons, RGCs rely on localized mRNA and protein synthesis to adapt rapidly to their environment, support axonal growth, and synaptic function (22, 23). By understanding the specific transcriptome of RGC axons, we could then better understand cellular processes within the axon related to neural development. To achieve this, RNA was isolated from both somatodendritic and axonal regions separately, and bulk RNA sequencing was performed using RNA from each region. In comparison to the somatodendritic compartment, RGC axons were enriched for many genes related to pathways associated with neuronal and axonal function, including neurotransmitter release, axon guidance, and voltage-gated potassium channels, including increased expression of genes such as SYN3 and CAMK4. Interestingly, we also found preferential localization of some transcripts associated with neurotransmitter receptor binding, including significant upregulation of genes such as GRIA4, suggesting perhaps a role in glutamate signaling onto axons associated with axonal growth and synaptic plasticity. Among the transcripts preferentially localized to RGC axons was also YBX1, a gene that functions in RNA metabolism and alternative splicing. Interestingly, YBX1 was also found to be preferentially localized to axons in a recent study analyzing hPSC-derived motor neurons (24, 25), suggesting a shared importance in axons across neuronal cell types. Conversely, the somatodendritic compartment was observed to have preferential localization of RNA transcripts associated with cell signaling and adhesion, including signaling factors such as BMP4 and SHH, with known roles in regulating retinal development (26–28), as well as CX3CL1, which serves as a primary axis for neuronal communication with microglia (29). We also found increased expression of adhesion genes including PLXNA1, known for a role in cell-cell adhesion, as well as the ECM component collagen type 2 (COL1A2), suggesting an increased need for cell adhesion in the somatodendritic compartment rather than the axonal compartment. This difference may reflect an increased level of maturation of the somatodendritic compartment, as RGCs are maintained in that chamber for multiple weeks while axons are dynamically growing over the timecourse of these experiments, perhaps decreasing their persistent reliance upon cell adhesion.

**Figure 2.**
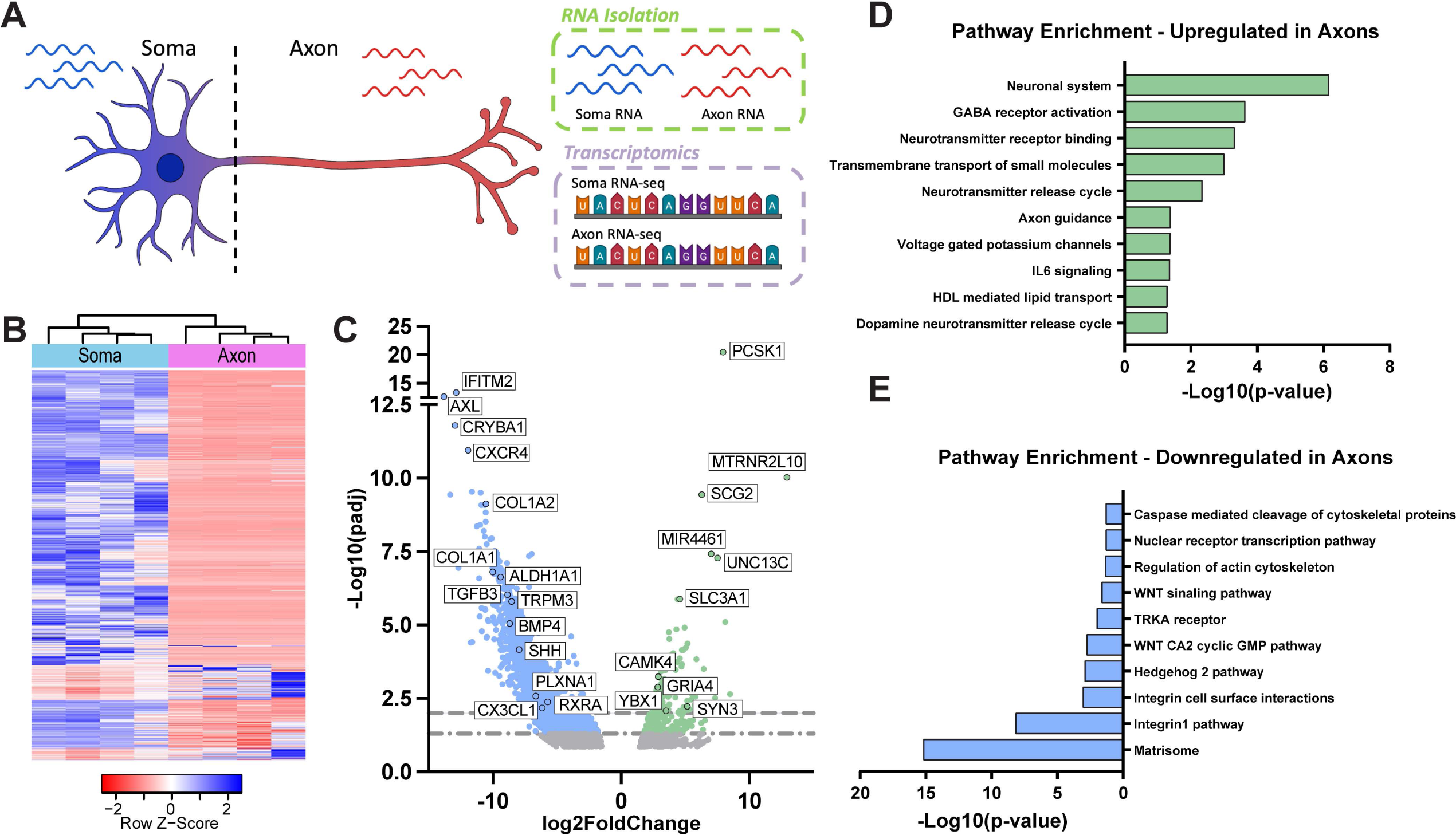
Transcriptional analysis shows RGC compartmentalization in microfluidic devices. (A) Schematic representation of hPSC-derived RGC neurite outgrowth on microfluidics devices and RNA isolation of both somatodendritic and axonal chamber separately. (B) Heatmap representing differentially expressed genes between soma and axon regions. Each row represents one sample, and each column represents one gene. The down- and up-regulated genes are red and blue colored, respectively. (C) Volcano plot of differentially expressed genes in the RGC axonal region compared to somatic region. (D-E) Gene Ontology (GO) and pathway terms from differential gene expression analysis of axonal and somatic regions.

### Microfluidic devices reveal axonal degenerative phenotypes in hPSC-RGC carrying the OPTN(E50K) glaucoma-associated mutation

In glaucoma, it is well-established that the primary site of injury occurs along the initial part of the axon, and degeneration of RGC compartments occurs by different mechanisms in the axonal and somatodendritic regions (16, 19). Therefore, RGC compartmentalization is an important feature to take into consideration when developing *in vitro* models for diseases like glaucoma (5, 15). Using hPSCs with an E50K mutation in the Optineurin (OPTN) gene, a leading cause of inherited forms of glaucoma (30, 31), we previously demonstrated neurite retraction in hPSC-RGCs carrying the OPTN(E50K) mutation when cultured in standard conditions (10, 11), yet it was not previously possible to confer these neurite retractions to either dendrites or axons due to considerable overlap in expression of markers associated with these respective neuronal compartments. Thus, we sought to leverage microfluidic platforms for the study of how the RGC somatodendritic and axonal compartments are affected by the OPTN(E50K) glaucoma associated mutation. RGCs were differentiated from hPSCs carrying the OPTN(E50K) mutation as well as their respective isogenic controls and were then seeded onto microfluidic platforms and maintained for 4 weeks to allow subsequent maturation (Figure 3A). Within these 4 weeks, we observed that both OPTN(E50K) RGCs and isogenic control RGCs were able to extend their axons toward the axonal chamber crossing the microgrooves, with MAP2-positive dendrites located only within the somatodendritic chamber, while SMI-312 positive axons extended into the axonal chamber (Figure 3B-C). We then analyzed morphological alterations in both somatodendritic and axonal regions individually (Figure 3D-K). Within the somatodendritic compartment, we observed a reduction in the soma size of OPTN(E50K)-RGCs compared to isogenic controls (Figure 3F), corroborating results we obtained previously without the use of microfluidic platforms (10, 11), while the number of primary MAP2-positive dendrites remained unchanged (Figure 3G), suggesting that previous neurite deficits may not have involved RGC dendrites. Conversely, within the axonal chamber, we observed decreased complexity of RGC axons, as measured by a reduction in the area occupied by the OPTN(E50K)-RGC axons (Figure 3J), as well as a reduction in axon extension and complexity measured by Sholl analysis (Figure 3K), compared to their respective isogenic controls. These results suggested that the OPTN(E50K) glaucoma-associated mutation resulted in a specific degeneration of RGC axons as opposed to dendrites, highlighting microfluidic platforms as powerful tools to model axonal degeneration that occurs in glaucoma.

**Figure 3.**
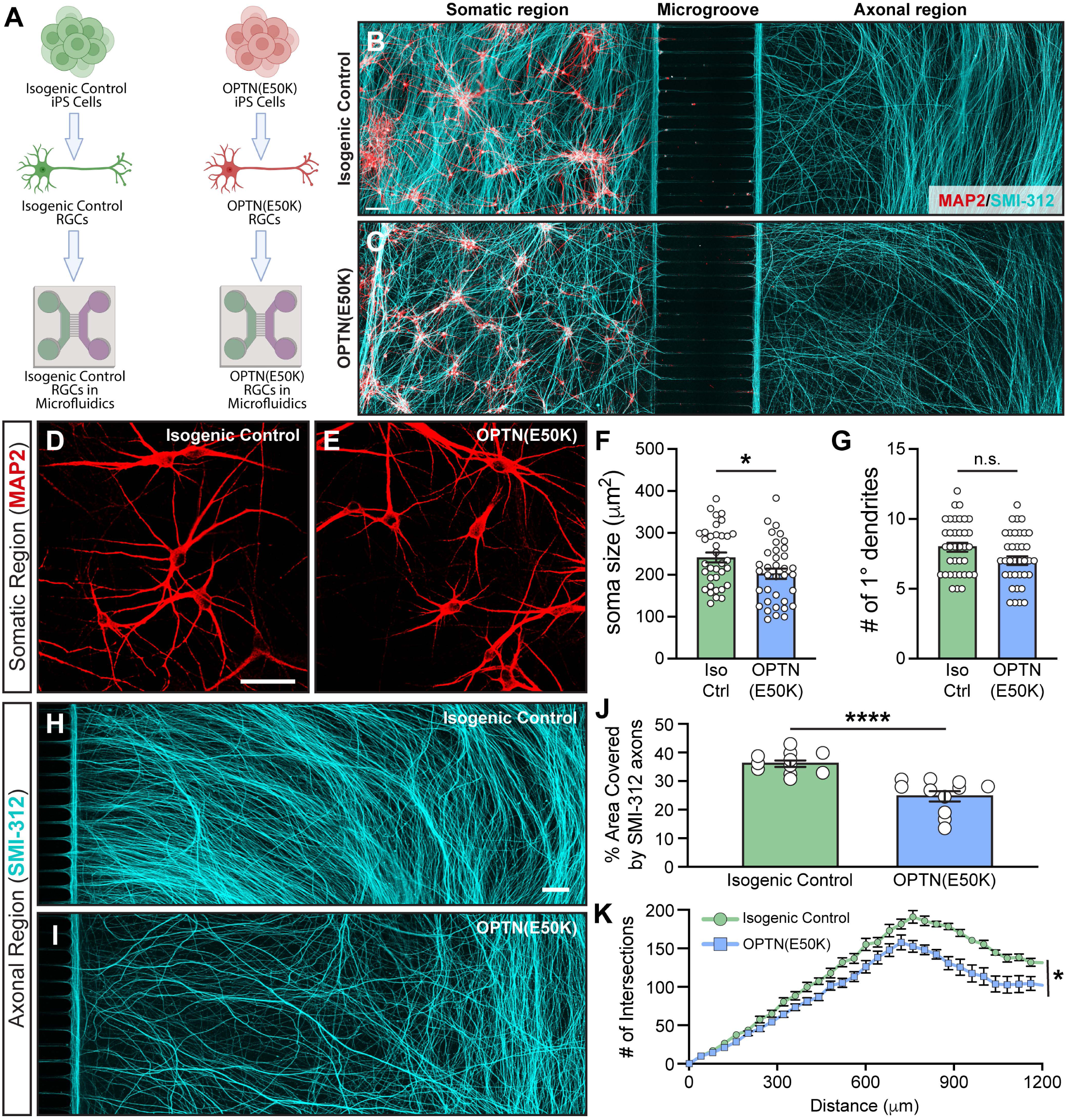
hPSC-RGC carrying the OPTN(E50K) glaucoma-associated mutation show morphological alterations in both somatodendritic and axonal compartments. (A) Schematic representation of RGCs differentiated from isogenic control or OPTN(E50K) iPSCs as well as seeding of RGCs from both cell lines on microfluidic devices. (B-C) Representative images of RGC somas (MAP2-positive dendrites) in the left chamber (somatic region) extending axons (SMI312-positive) through the contralateral chamber (axonal region), differentiated from isogenic control or OPTN(E50K) hPSCs, respectively. Scale bar: 100 μm. (D-E) High magnification images showing MAP2-positive dendrites for both isogenic control and OPTN(E50K) RGCs, respectively, in the somatic region of microfluidic platforms. Scale bar: 50 μm. (F-G) Quantification of soma size and the number of primary MAP2-positive dendrites in isogenic control and OPTN(E50K) hPSC-RGCs (n=36 for each WT and E50K from 3 biological replicates; *t*-test, *p*=0.027 for F and *p*=0.963 for G) show a significant decrease in the soma size of OPTN(E50K) RGCs compared to controls. (H-I) Representative images of SMI-312 positive axons in the axonal region. Scale bar: 100 μm. (J-K) Quantification of axonal area and Sholl analysis in isogenic control and OPTN(E50K) hPSC-RGCs (J: n=11 images for each WT and E50K from 3 biological replicates; K: n=11 images for each WT and E50K from 3 biological replicates; *t*-test, *p<0.05, ****p<0.0001). Error bars represents mean values ±LS.E.M.

Among axonal deficits associated with glaucoma, previous studies have also shown that axonal transport is compromised within RGC axons (32) and as such, we explored whether axonal transport deficits would be observed in RGC axons grown in microfluidic platforms from isogenic OPTN(E50K) RGCs compared to unaffected controls. To accomplish this, RGCs were incubated with MitoTracker to label mitochondria (Figure 4), which are readily transported both anterogradely and retrogradely along the length of the axon. Mitochondria also play a critical role in the long-distance projections of RGC axons, as increased mitochondrial density is required in the unmyelinated optic nerve to fulfill metabolic function in RGC axons (33, 34), and the OPTN(E50K) mutation is known to result in mitochondrial deficits in RGCs and astrocytes (20, 35). Initially, the labeling of mitochondria in RGC axons with MitoTracker demonstrated a significant increase in the number of mitochondria within OPTN(E50K) axons, associated with significant decreases in OPTN(E50K) mitochondrial length and shape or aspect ratio (Figure 4B-D, respectively). Moreover, since OPTN serves as a critical mitophagy receptor to aid in the removal or recycling of damaged mitochondria (36, 37), we then examined whether the OPTN(E50K) mutation disrupts mitochondrial transport in the axonal compartment. The movement of MitoTracker-labeled mitochondria was recorded over a period of 5 minutes, as represented in kymographs (Figure 4E-F). When compared to isogenic controls, an increased number of stationary mitochondria were observed in the axons of OPTN(E50K) RGCs (Figure 4F), which was found to occur at the expense of retrograde transport, while anterograde transport remained unchanged (Figure 4G). Furthermore, while no alterations were observed in the velocity of transported mitochondria (Figure 4H), retrogradely transported mitochondria were observed to be transported significantly further distances in the axons of OPTN(E50K) RGCs (Figure 4I). Collectively, these results demonstrated that the use of microfluidic platforms to isolate RGC axons enabled the identification of mitochondrial alterations in RGC axons as well as changes in axonal transport, suggesting that the OPTN(E50K) mutation negatively affected axonal transport in RGCs with this gene mutation.

**Figure 4.**
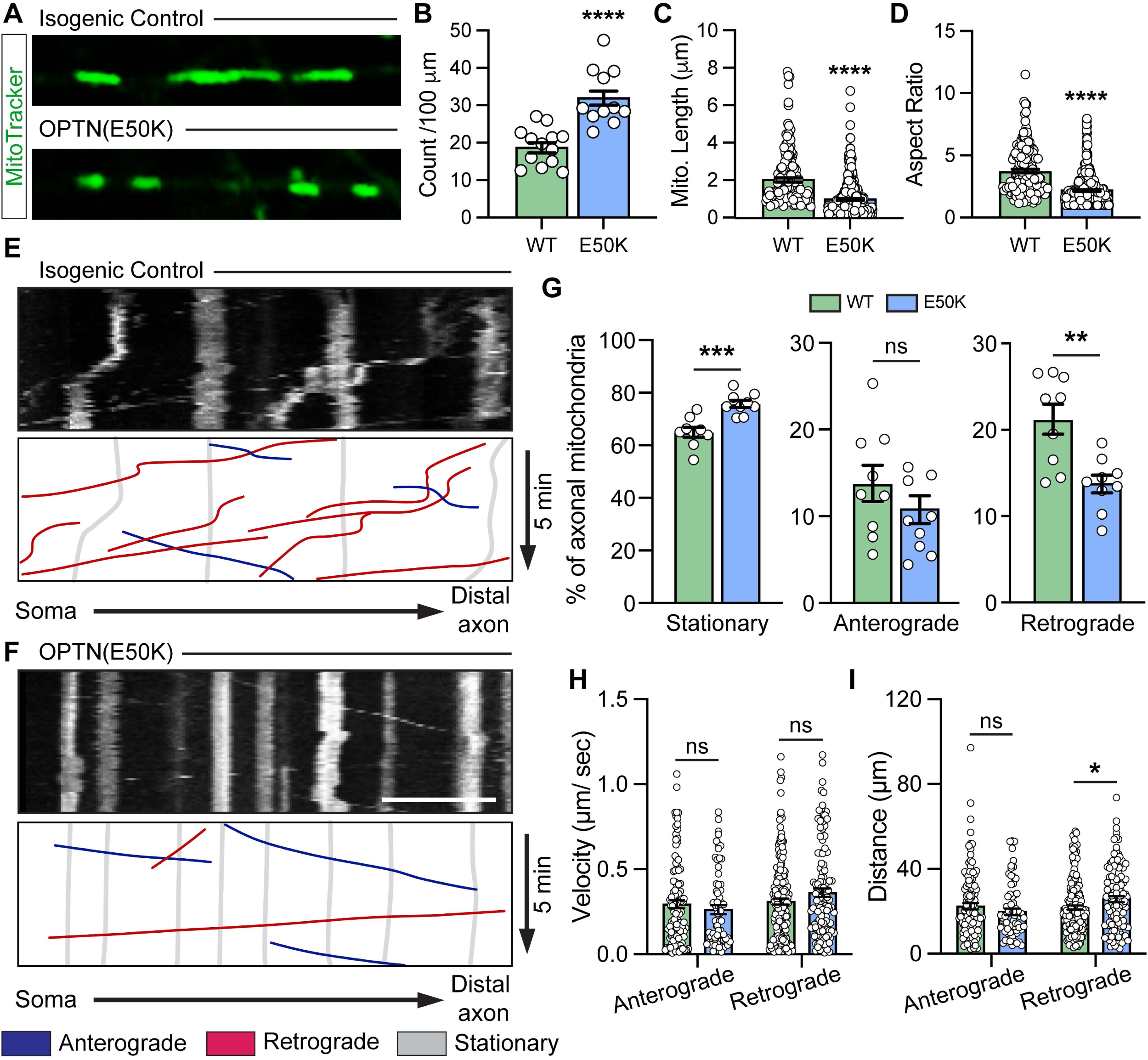
Mitochondria deficits in the axons of OPTN(E50K) mutant hPSC-RGC. (A) Representative images demonstrating isogenic control and OPTN(50K) RGCs stained with MitoTracker Green. Scale bar: 10 μm. (B-D) Quantification of mitochondria number, length and aspect ratio in both isogenic control and OPTN(E50K) hPSC-derived RGCs (B: n=13 SMI-312 axon images for each WT and E50K from 3 biological replicates; C and D: n=170 for WT and n=349 for E50K mitochondria counts from 13 SMI-312 axon images from 3 biological replicates). (E-F) Representative images of mitochondria movement kymographs of 5 min recording. X-axis represents distance from the soma (proximal axon) to distal axon. Y-axis represents the recording time (5 min). Scale bar: 5 μm. (G-I) Quantification of each mitochondria movement separately, velocity, and distance in isogenic control and OPTN(E50K) RGCs (F: n=9 SMI-312 axon recordings for each WT and E50K from 3 biological replicates; G and H: n=112 for WT anterograde, n=69 for E50K anterograde, n=176 for WT retrograde, and n=118 for E50K retrograde mitochondria counts from 9 SMI-312 axon recordings with 3 biological replicates; *t*-test, *p<0.05, ns>0.05). *t*-test, *p<0.05, **p<0.01, ***p<0.001, p<0.0001, ns>0.05 Error bars represents mean values ±LS.E.M.

Given the morphological and functional changes we observed in RGC axons between OPTN(E50K) and isogenic control cells, we next sought to further characterize the transcriptional profile of RGC axons in these two types of samples using axon-sequencing techniques (38). Both OPTN(E50K) and isogenic control RGCs were grown in microfluidic platforms to isolate their axons into the distinct axonal compartment, and RNA was isolated from the axons of both cell types (Figure 5A). Axonal RNA sequencing was performed to determine differentially expressed genes in the axonal compartment of OPTN(E50K) RGCs compared to isogenic control RGC axons. Interestingly, among the significantly differentially expressed genes between isogenic control and OPTN(E50K) axons (Figure 5B), we observed an increase in the expression of genes such as MAPKBP1, TRIM46, and CDK5 that have been previously described to play a role in the maintenance of the structural framework of axons (39–42). Additionally, we also observed a significant increase in the expression of genes such as PRMT8 and CACNA1B, which contribute to intracellular signaling and synaptic activity (43, 44), as well as NINJ1, which is involved in the maintenance of axonal integrity and axonal regeneration responses (45). Conversely, we found a significant decrease in OPTN(E50K) axons of genes such as PLXNC1, FREM3, and SFRP1, which have been shown to have roles in axonal pathfinding and guidance (46, 47), as well as genes such as MAP7D3 and GPNMB which have been associated with axonal structure and maintenance (48, 49). Therefore, the reduced expression of these genes may contribute to the degenerative axonal phenotype observed in OPTN(E50K) RGCs compared to isogenic controls.

**Figure 5.**
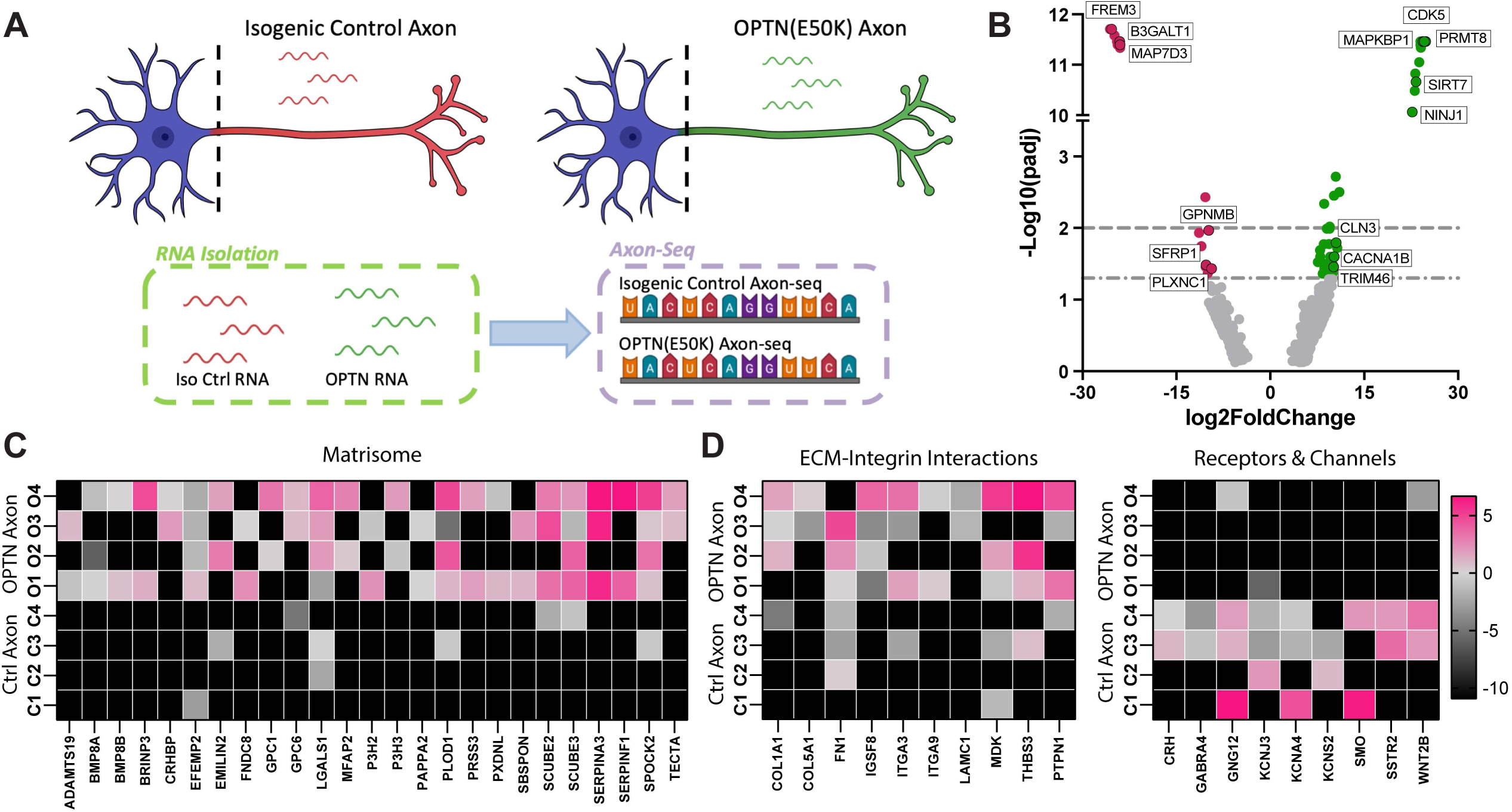
Axon-sequencing of isogenic control and OPTN(E50K) hPSC-derived RGCs. (A) Schematic representation of isogenic control and OPTN(E50K) hPSC-derived RGCs seeded in microfluidic chips highlighting the axonal region from where RNA was extracted to be used in axon-sequencing. (B) Volcano plot representing log2(fold change) gene expression between the axonal compartment of OPTN(E50K) RGCs *versus* isogenic controls, plotted with respect to - log10(padj). (C-F) Heatmaps of genes expressed within matrisome (C), extracellular matrix (ECM)-integrin interactions (D) as well as receptors and channels (E) signaling pathways represented based on normalized counts. The down- and up-regulated genes are black and fuchsia colored, respectively.

Through pathway enrichment analysis, we identified the matrisome and extracellular matrix (ECM) as the most upregulated signaling pathways observed in the axonal region of OPTN(E50K) RGCs (Figure 5C and D). Recent studies have shown the crucial role of ECM and matrisome during retinal development, including their contribution to axonal growth and guidance (50, 51). The alterations observed in these signaling pathways in the axons of OPTN(E50K) glaucoma-associated RGCs may contribute to axonal degeneration as the dysregulation of the ECM components has been identified in several disease conditions (52), including glaucomatous neurodegeneration (53). On the other hand, genes associated to receptors and ion channels were found among the most downregulated in the axonal compartment of OPTN(E50K) RGCs (Figure 5F). Voltage gated potassium channels are crucial for RGC development, neurite outgrowth, axon guidance and function due to their crucial role in the generation and propagation of action potential (54, 55). Therefore, the reduced expression of potassium channels in the axonal compartment of OPTN(E50K) RGCs may affect their function and contribute to glaucoma-associated axonal degeneration. Interestingly, the downregulation of GABRA41, a member of the GABA-A receptor gene family, may also be involved in axonal degeneration RGCs since this receptor has been shown to be expressed by RGCs (56) and the downregulation of GABA receptors linked to degraded neural specificity the visual cortex of glaucoma patients (57). Altogether, these results highlight axon-sequencing as an important tool to identify differentially expressed genes and signaling pathways that specifically affect the degeneration of RGC axons in glaucoma models.

### Reactive astrocytes located in the proximal axonal compartment of RGCs promote a reduction in axonal extension in distal axons

Across many underlying causes of glaucoma, the induction of glial reactivity in the optic nerve head is strongly associated with the phenotypes observed in glaucoma patient eyes (58). The focal nature of glial reactivity and localized injury to RGC axons requires greater attention to the highly compartmentalized nature of glaucomatous injury. Therefore, microfluidic platforms with three chambers were established containing RGCs and astrocytes differentiated from hPSCs, with astrocytes seeded along the proximal axonal region of RGCs to specifically study the role of reactive astrocytes over RGC axons (Figure 6). Astrocytes were induced and maintained in a reactive state through incubation with TNFα, IL-1α, and C1q (59–61). Our results demonstrated that RGCs extended their axons through both of the two axonal chambers (SMI-312 positive axons), while somas and dendrites remained uniquely in the soma chamber (MAP2 positive dendrites) (Figure 6A-B). Moreover, when the astrocytes plated in the proximal axonal chamber were induced to a reactive state, neurotoxic effects were observed in both the somatodendritic region and distal axonal compartment of the RGCs. Reactive astrocytes located along the proximal axonal compartment of RGCs promoted a reduction in the number of primary MAP2-positive dendrites and soma size (Figure 6C-F), compared to the effect promoted by control astrocytes. Importantly, reactive astrocytes induced a reduction in the area occupied by the axons in the distal axonal compartment, as well as a reduction in axonal extension and complexity (Figure 6G-J). These results highlighted these three-chamber microfluidic platforms as a powerful model to study the effect of reactive astrocytes in an appropriately spatially oriented manner along RGC axons to examine glial neuroinflammatory roles to RGC neurodegenerative features.

**Figure 6.**
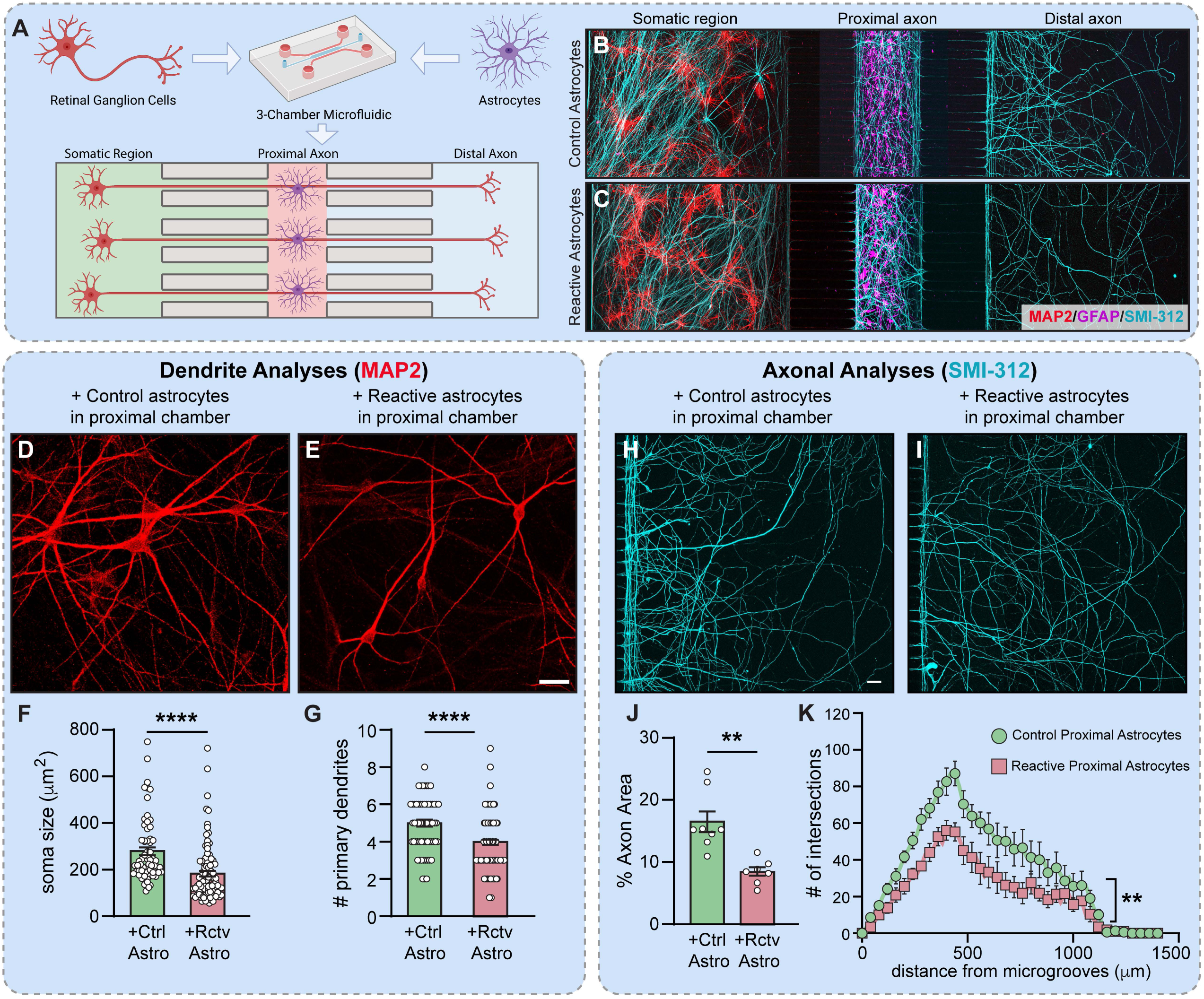
Reactive astrocytes located in the proximal axonal compartment promote degenerative phenotypes in RGC distal axons. (A) Schematic representation of hPSC-derived RGC seeded in the somatic region and extending axons throughout the proximal axon and distal axon regions with hPSC-derived astrocytes seeded in the proximal chamber in close contact with the proximal axonal compartment of RGCs. (B-C) Representative images of RGC somas (MAP2-positive) in the left chamber (soma region) extending axons (SMI312-positive) through the second and third chambers (proximal and distal axon, respectively). Astrocytes cultured in the proximal chamber (GFAP-positive) mimicking the location of glia relative to RGCs in the optic nerve. (D-E) MAP2 staining in the soma region showing RGC morphology after culture with control or reactive astrocytes in the proximal chamber. (F-G) Quantification of soma size and the number of primary neurites in RGCs exposed to the effect of control or reactive astrocytes located in the proximal chamber (n=66 and 98 RGCs, cultured with control and reactive astrocytes, respectively, from 4 biological replicates). (H-I) Representative images of SMI-312 labeled axons in the distal chamber, as well as Sholl analysis representation. (J-K) Sholl analysis and quantification of the area occupied by the RGC axons after culture with control or reactive astrocytes located in the proximal chamber (n=4 biological replicated using a total of 2 images per condition). Data represent mean values ± S.E.M. *pL<L0.05, **pL<L0.01 and ***pL<L0.001 *vs.* control astrocytes, two-tailed unpaired Student’s t test. Scale bar: 100 μm.

## DISCUSSION

Axons are crucial components of neurons, serving as the primary transmission lines for electrical signals across the nervous system. In RGCs, axons play a particularly vital role, with these axons extending from the retina to the brain, forming the optic nerve and transmitting visual information to processing centers such as the lateral geniculate nucleus and the visual cortex (1, 2). The integrity and function of axons in RGCs are essential for proper vision, as they facilitate the communication required for interpreting visual stimuli. Damage or degeneration of these axons, as seen in conditions like glaucoma, can lead to vision loss or blindness (62). Thus, the axon’s role in transmitting neural signals is fundamental to the overall function of the visual system, yet the proper compartmentalization of hPSC-derived RGCs has received relatively little attention to date, serving as the justification for the studies outlined within this manuscript.

Our initial efforts focused upon the identification and localization of markers traditionally associated with the somatodendritic and axonal compartments of neurons, particularly MAP2 and SMI-312, respectively. Within traditional 2D cultures of purified RGCs, we observed robust expression of MAP2 and SMI-312, although some of this expression overlapped, suggesting an incomplete degree of maturation of these neurons. Conversely, when grown in microfluidic platforms, we observed a more complete compartmentalization of RGCs, including a lack of any overlap between MAP2 and SMI-312, as well as more extensive dendritic branching, suggesting that RGCs achieved a greater degree of maturation within microfluidic platforms. This could potentially be explained by the gradient of BDNF created by supplementing the distal microfluidic chamber with higher levels of BDNF, resulting in more polarized neurons due to long distance axonal outgrowth. At the same time, we recognize that other possible contributing factors exist, including the surface of microfluidic chips consisting of a cyclic olefin copolymer plastic whereas RGCs grown in traditional 2D cultures were grown on glass coverslips, although both were coated with laminin prior to seeding of RGCs.

Perhaps most importantly, our finding that RGC axons could be uniquely isolated into distal chambers of the microfluidic chip provides novel opportunities for the investigation of axon-specific phenotypes, both from a developmental standpoint as well as in disease states. Initially, our observation that MAP2 and SMI-312 could be reliably used to identify the somatodendritic and axonal compartments, respectively, when RGCs were grown in microfluidic compartments provides novel opportunities to identify unique features to each of these neuronal compartments, as well as disease-related phenotypes specific to each compartment as well. Among the axonal phenotypes that have been previously associated with the optic nerve in glaucoma, deficits in axonal transport are often cited as a defining characteristic (32, 62). As such, we sought to explore deficits in axonal transport associated with glaucoma in this human RGC microfluidic model by labeling mitochondria with MitoTracker and observing the motility of mitochondria up or down the axon within the axonal chamber of the microfluidic platform. The transport of mitochondria along the axon was chosen to examine deficits in axonal transport due to 1) simplicity to label mitochondria with the MitoTracker reagent, 2) ability to perform live-cell imaging to observe real-time mitochondrial movement along axons, and 3) the deficient transport of mitochondria has been previously observed in models of glaucoma, and 4) the OPTN(E50K) mutation has been previously associated with dysfunction of autophagy and by extension, mitophagy. We initially observed that mitochondria in OPTN(E50K) axons were smaller and more numerous than in isogenic control RGC axons, with a rounder appearance (aspect ratio), suggestive of mitochondrial fission. Interestingly, prior studies have demonstrated that dysregulated mitochondrial fission contributes to axonal and neuronal damage in glaucoma and other optic neuropathies, which has been associated with clearance by mitophagy (63, 64). In the context of the OPTN(E50K) mutation, it is possible that autophagic (and mitophagic) dysfunction impedes this clearance of damaged mitochondria, leading to their retention and associated with overall mitochondrial dysfunction.

Additionally, we sought to more specifically examine changes in mitochondrial axonal transport, where we observed an increase in the percentage of stationary mitochondria in OPTN(E50K) RGC axons, which came at the expense of retrograde transport but not anterograde transport. Prior studies have demonstrated that impaired retrograde transport prevents the delivery of neurotrophic factors such as BDNF to the soma (65), which is a known retrograde-derived survival factor for RGCs (66). The accumulation of damaged organelles such as mitochondria in axons also exacerbates stress responses and promotes neurodegeneration (67). While some previous studies have demonstrated anterograde transport deficits in other models of glaucoma, this is more typically associated with later stages of the degenerative process, contributing to energy shortages in distal axons and synaptic degeneration. Conversely, other studies have shown that early stages of glaucoma often show retrograde transport impairment, affecting processes such as neurotrophic signaling, making RGCs more vulnerable to stress (68). As *in vitro* stem cell models mimic developmental principles in a dish, it stands to reason that earlier axonal phenotypes such as retrograde transport dysfunction would be primarily observed within these models.

With the ability to isolate and analyze the axonal compartment apart from the somatodendritic compartment, we were also able to assess the transcriptional profile of RGC axons. It has been known that certain RNA transcripts are preferentially localized to the axonal compartment, where local RNA translation plays a critical role in supporting the dynamic functions of the axon by enabling localized protein synthesis and allowing axons to rapidly respond to changes in their environment, such as synaptic activity, injury, or growth signals, without needing to rely on protein transport from the cell body (22, 23). Recent studies have pursued these types of analyses in other classes of neurons such as motor neurons (38), but it has not been known to date how the transcriptional profile of RGC axons differs from their somatodendritic counterpart as well as in comparison to axons in a diseased state. To pursue these analyses, we performed Axon-seq methods to assess the transcriptional profile of RGC axons. We observed the preferential localization of some transcripts associated with axonal functions within the axonal compartment, including pathways such as axon guidance and neurotransmitter release. Interestingly, pathway enrichment analyses also demonstrated increased expression of genes associated with neurotransmitter receptor binding and GABA receptor activation, both of which would typically be expected within RGC dendrites, suggesting a potential role for these neurotransmitters to regulate axonal functions such as modulation of axonal growth, plasticity, and even local protein translation (56). Additionally, the identification of transcripts in the axon associated with the dopamine neurotransmitter release cycle is reminiscent of certain subtypes of RGCs. While many RGCs are known to be glutamatergic, recent studies have suggested that dopamine from RGCs during development may regulate retinal angiogenesis, or it may represent a subtype of RGC within this population (69, 70). It is also important to consider that the somatodendritic chamber of the microfluidic platforms was seeded with RGCs that were highly enriched from cultures following MACS purification, but this does not necessarily mean that other cell types were not present. Indeed, over the course of weeks required for experimental analyses, it is possible that an initial small population of retinal progenitor cells remained within the sorted RGC population. While insignificant in number at the point of purification, their ability to grow and divide over the next few weeks combined with the post-mitotic nature of RGCs provides the possibility that the transcripts observed in the somatodendritic chamber of microfluidic platforms was largely due to RGC somas, but could also contain some proliferative retinal progenitors or other retinal neurons derived from them. Given this possibility, it is likely best to focus primarily upon the transcriptional analysis of genes and pathways preferentially expressed in RGC axons, as the axonal chamber of microfluidic platforms remained a pure population of RGC axons in the absence of any other cells or cellular processes.

When assessing Axon-seq results comparing isogenic control and OPTN(E50K) axons, we observed clear differences between the two genotypes. Initially, it was apparent that more transcripts were more highly expressed in OPTN(E50K) axons compared to isogenic control axons than the inverse situation. While one would expect the total number of transcripts detected to be limited in the axonal compartment overall, we found that there were more transcripts that were significantly higher in expression in OPTN(E50K) axons compared to isogenic control axons, whereas the number of transcripts higher in expression in isogenic control axons compared to OPTN(E50K) axons was more limited. Among these differentially expressed genes, many of those more highly expressed in isogenic control axons had functions related to axonal and neuronal function, including Microtubule-Associated Protein 7 Domain-Containing Protein 3 (MAP7D3), which regulates the stability and organization of microtubules (71), or a variety of potassium channels, including KCNA4 and KCNS2 which are pore-forming subunits of voltage-gated potassium channels and whose dysregulation is associated with hyperexcitabilty. Interestingly, our previous studies have also demonstrated a hyperexcitable phenotype in OPTN(E50K) RGCs (11). Conversely, and perhaps more interestingly, was the fact that many of the transcripts expressed more highly in OPTN(E50K) axons were associated with the extracellular matrix and matrisome. Given the likely changes in the extracellular environment surrounding OPTN(E50K) axons, it is possible that these changes would not only affect axonal function, but would also likely influence other neighboring cells, such as glia and vascular cells. In future studies, it would be of interest to explore how other cell types would respond to this altered extracellular environment.

Given the ability to induce long-distance RGC axonal projects into separate chambers, our final results included the use of microfluidic platforms with three chambers to further subdivide the axons into “proximal” and “distal” axonal compartments. Since the primary insult in glaucoma occurs to axons as they traverse the lamina cribrosa and exit the eye into the brain (72), the proximal region of the axon is of particular significance. While the current studies do not address the mechanical forces resulting in axonal strain and compression at the optic nerve head, the inflammatory response from glia at that region is also of great importance, resulting in a localized inflammatory response in the proximal region of RGC axons (58). As we have recently demonstrated that reactive, pro-inflammatory astrocytes result in neurodegenerative features in RGCs as a whole (59), the introduction of hPSC-derived astrocytes within the middle chamber to simulate the proximal region of the axon led to morphological changes in RGCs when astrocytes were stimulated to a pro-inflammatory state. Interestingly, unlike OPTN(E50K) RGCs as described in Figure 3 where in the absence of glia the E50K mutation resulted only in significant changes in the axonal compartment, the introduction of pro-inflammatory astrocytes along the proximal region of the axon led to both distal axonal retraction as well as a reduction in dendrites. These bi-directional effects of pro-inflammatory astrocytes in the proximal axonal region are of great interest, and future studies will be needed to determine the consequences of these effects, including perhaps the further advancement of these models to incorporate other cell types such as microglia and/or vascular cells.

While the current studies represent a noteworthy advance in the development of an *in vitro* platform that better mimics RGC compartmentalization, future studies will also be needed to address other shortcomings that remain. The division of the axon into proximal and distal regions as described in Figure 6 is an important advance in the study of spatial orientation of cell types along the optic nerve, future studies will certainly be needed to better develop these platforms into something that more closely mimics the biomechanics of the lamina cribrosa (73). Another interesting feature of the optic nerve is its myelination pattern, where RGC axons are unmyelinated prior to the optic nerve head, but become myelinated after they exit the eye (74). The current system was devoid of any oligodendrocytes, but previous studies have demonstrated the ability to derive oligodendrocytes from hPSCs and it would be of considerable interest in future studies to leverage these microfluidic platforms to explore RGC axonal myelination. Finally, although RGCs within these microfluidic platform extended lengthy axons, the current system does not account for post-synaptic targets in the brain (75). In future studies, it would be of considerable interest to introduce target cell types of the thalamus into the distal axonal chamber for studies of synaptic connectivity and retrograde communication from thalamus to RGCs.

## MATERIALS AND METHODS

### Maintenance and expansion of hPSCs

Details about the hPSC lines used in this study, as well as the procedures regarding hPSC maintenance and expansion can be found in the Supplemental Information. Briefly, hPSCs were maintained as previously described (26, 76), in mTeSR1 medium on either a Matrigel- or Geltrex-coated substrate, and expanded through the use of ReLeSR reagent.

### Retinal organoid differentiation and RGCs isolation

hPSCs were differentiated into RGCs using previously published protocols (11, 76–78), and as detailed in the Supplemental Information.

### Astrocyte differentiation and induction of a reactive state

hPSCs were differentiated to an astrocytic lineage according to previously established protocols (20, 21, 79), and as detailed in the Supplemental Information. To induce a reactive phenotype, hPSC-derived astrocytes were incubated with TNFα (30 ng/mL), IL-1α (3 ng/mL), and C1q (400 ng/mL), as previously described (20, 21).

### Cell seeding and maintenance of microfluidic devices

The microfluidic devices were coated and handled following manufacturer’s instructions and as previously described (80). 2-compartment XonaChips (XC450) and 3-compartment XonaChips (XC-T500) from Xona microfluidics^®^ (North Carolina, USA) were used in this study. Detailed procedures are provided in the Supplemental Information.

### Immunocytochemistry

hPSC-derived RGCs seeded on coverslips or microfluidic devices were fixed with 4% paraformaldehyde at the indicated timepoints, and immunostained as previously described (20, 78, 81). Detailed procedures are provided in the Supplemental Information.

### Dendritic and axonal quantification

For morphological analysis and measurements, MAP2 and SMI-312 fluorescent images were analyzed using the ImageJ (Fiji) software. Moreover, Imaris Image Analysis Software was used to determine the co-localization of MAP2-positive dendrites and SMI-312 axons in standard and microfluidic cultures. Detailed procedures are provided in the Supplemental Information.

### Mitochondria transportation recording

Mitochondria length and movement were determined in hPSC-derived RGCs using MitoTracker Green, after 4 weeks of maturation, as previously described (82), with modifications, and as detailed in the Supplemental Information.

### RNA isolation and mRNA sequencing

Detailed procedures related to RNA isolation from microfluidic platforms as well as to mRNA sequencing and respective analysis are provided in the Supplemental Information.

### Statistical analysis

Detailed description of the statistical analyses performed for each result is available in the supplemental information.

## ACKNOWLEDGMENTS

We would like to thank the Indiana University Center for Medical Genomics for assistance with RNA-seq analyses. Grant support was provided by the National Eye Institute (R01EY033022 and U24EY033269 to JSM), the BrightFocus Foundation (G2022014S to JSM), the Gilbert Family Foundation (923016 to JSM), the Glaucoma Research Foundation (to JSM), and the Indiana Department of Health Spinal Cord and Brain Injury Research Fund (26343 to JSM). Support for this project was also provided by the BrightFocus Postdoctoral Fellowship (G2022003F to CG) and the Shaffer Award from the Glaucoma Research Foundation (to CG), as well as a Cagiantas scholarship from the Indiana University School of Medicine (JH).

## Notes

### Competing Interest Statement

The authors have declared no competing interest.

## REFERENCES

1. M. C. Crair, C. A. Mason, Reconnecting Eye to Brain. J Neurosci 36, 10707–10722 (2016).

2. L. Erskine, E. Herrera, Connecting the retina to the brain. ASN Neuro 6 (2014).

3. L. Conforti, R. Adalbert, M. P. Coleman, Neuronal death: where does the end begin? Trends Neurosci 30, 159–166 (2007).

4. D. Y. Yu et al., Retinal ganglion cells: Energetics, compartmentation, axonal transport, cytoskeletons and vulnerability. Prog Retin Eye Res 36, 217–246 (2013).

5. A. V. Whitmore, R. T. Libby, S. W. John, Glaucoma: thinking in new ways-a role for autonomous axonal self-destruction and other compartmentalised processes? Prog Retin Eye Res 24, 639–662 (2005).

6. A. Artero-Castro et al., Glaucoma as a Neurodegenerative Disease Caused by Intrinsic Vulnerability Factors. Prog Neurobiol 193, 101817 (2020).

7. J. R. Soucy et al., Retinal ganglion cell repopulation for vision restoration in optic neuropathy: a roadmap from the RReSTORe Consortium. Mol Neurodegener 18, 64 (2023).

8. F. Yuan, M. Wang, K. Jin, M. Xiang, Advances in Regeneration of Retinal Ganglion Cells and Optic Nerves. Int J Mol Sci 22 (2021).

9. C. M. Fligor et al., Three-Dimensional Retinal Organoids Facilitate the Investigation of Retinal Ganglion Cell Development, Organization and Neurite Outgrowth from Human Pluripotent Stem Cells. Sci Rep 8, 14520 (2018).

10. K. C. Huang et al., Acquisition of neurodegenerative features in isogenic OPTN(E50K) human stem cell-derived retinal ganglion cells associated with autophagy disruption and mTORC1 signaling reduction. Acta Neuropathol Commun 12, 164 (2024).

11. K. B. VanderWall et al., Retinal Ganglion Cells With a Glaucoma OPTN(E50K) Mutation Exhibit Neurodegenerative Phenotypes when Derived from Three-Dimensional Retinal Organoids. Stem Cell Reports 15, 52–66 (2020).

12. M. L. Risner et al., Intrinsic Morphologic and Physiologic Development of Human Derived Retinal Ganglion Cells In Vitro. Transl Vis Sci Technol 10, 1 (2021).

13. V. M. Sluch et al., Enhanced Stem Cell Differentiation and Immunopurification of Genome Engineered Human Retinal Ganglion Cells. Stem Cells Transl Med 6, 1972–1986 (2017).

14. P. Teotia et al., Modeling Glaucoma: Retinal Ganglion Cells Generated from Induced Pluripotent Stem Cells of Patients with SIX6 Risk Allele Show Developmental Abnormalities. Stem Cells 35, 2239–2252 (2017).

15. A. Donato, K. Kagias, Y. Zhang, M. A. Hilliard, Neuronal sub-compartmentalization: a strategy to optimize neuronal function. Biol Rev Camb Philos Soc 94, 1023–1037 (2019).

16. S. B. Syc-Mazurek, R. T. Libby, Axon injury signaling and compartmentalized injury response in glaucoma. Prog Retin Eye Res 73, 100769 (2019).

17. A. M. Taylor, N. L. Jeon, Microfluidic and compartmentalized platforms for neurobiological research. Crit Rev Biomed Eng 39, 185–200 (2011).

18. E. Neto et al., Compartmentalized Microfluidic Platforms: The Unrivaled Breakthrough of In Vitro Tools for Neurobiological Research. J Neurosci 36, 11573–11584 (2016).

19. K. C. Huang, C. Gomes, J. S. Meyer, Retinal Ganglion Cells in a Dish: Current Strategies and Recommended Best Practices for Effective In Vitro Modeling of Development and Disease. Handb Exp Pharmacol 281, 83–102 (2023).

20. C. Gomes et al., Astrocytes modulate neurodegenerative phenotypes associated with glaucoma in OPTN(E50K) human stem cell-derived retinal ganglion cells. Stem Cell Reports 17, 1636–1649 (2022).

21. K. B. VanderWall et al., Astrocytes Regulate the Development and Maturation of Retinal Ganglion Cells Derived from Human Pluripotent Stem Cells. Stem Cell Reports 12, 201–212 (2019).

22. C. J. Costa, D. E. Willis, To the end of the line: Axonal mRNA transport and local translation in health and neurodegenerative disease. Dev Neurobiol 78, 209–220 (2018).

23. I. Dalla Costa et al., The functional organization of axonal mRNA transport and translation. Nat Rev Neurosci 22, 77–91 (2021).

24. J. Feng et al., Regulation of neurotransmitter release by synapsin III. J Neurosci 22, 4372–4380 (2002).

25. R. R. Ji et al., Ca2+/calmodulin-dependent protein kinase type IV in dorsal root ganglion: colocalization with peptides, axonal transport and effect of axotomy. Brain Res 721, 167–173 (1996).

26. J. Harkin et al., A highly reproducible and efficient method for retinal organoid differentiation from human pluripotent stem cells. Proc Natl Acad Sci U S A 121, e2317285121 (2024).

27. J. Huang, Y. Liu, A. Oltean, D. C. Beebe, Bmp4 from the optic vesicle specifies murine retina formation. Dev Biol 402, 119–126 (2015).

28. L. Zhao, H. Saitsu, X. Sun, K. Shiota, M. Ishibashi, Sonic hedgehog is involved in formation of the ventral optic cup by limiting Bmp4 expression to the dorsal domain. Mech Dev 127, 62–72 (2010).

29. J. H. Zhao, et al., The effect of CX3CL1/CX3CR1 signal axis on microglia in central nervous system diseases. J Neurorestoratology 11 (2023).

30. Y. Minegishi, M. Nakayama, D. Iejima, K. Kawase, T. Iwata, Significance of optineurin mutations in glaucoma and other diseases. Prog Retin Eye Res 55, 149–181 (2016).

31. Q. Meng et al., Overexpressed mutant optineurin(E50K) induces retinal ganglion cells apoptosis via the mitochondrial pathway. Mol Biol Rep 39, 5867–5873 (2012).

32. E. T. Fahy, V. Chrysostomou, J. G. Crowston, Mini-Review: Impaired Axonal Transport and Glaucoma. Curr Eye Res 41, 273–283 (2016).

33. T. Tsuji, T. Murase, Y. Konishi, M. Inatani, Optic Nerve Injury Enhanced Mitochondrial Fission and Increased Mitochondrial Density without Altering the Uniform Mitochondrial Distribution in the Unmyelinated Axons of Retinal Ganglion Cells in a Mouse Model. Int J Mol Sci 24 (2023).

34. N. A. Muench et al., The Influence of Mitochondrial Dynamics and Function on Retinal Ganglion Cell Susceptibility in Optic Nerve Disease. Cells 10 (2021).

35. M. Surma et al., Enhanced mitochondrial biogenesis promotes neuroprotection in human pluripotent stem cell derived retinal ganglion cells. Commun Biol 6, 218 (2023).

36. Y. Jeong et al., Glaucoma-associated Optineurin mutations increase transmitophagy in a vertebrate optic nerve. bioRxiv 10.1101/2023.05.26.542507 (2023).

37. K. Yamano et al., Optineurin provides a mitophagy contact site for TBK1 activation. Embo j 43, 754–779 (2024).

38. J. Nijssen, J. Aguila, R. Hoogstraaten, N. Kee, E. Hedlund, Axon-Seq Decodes the Motor Axon Transcriptome and Its Modulation in Response to ALS. Stem Cell Reports 11, 1565–1578 (2018).

39. P. Grant, P. Sharma, H. C. Pant, Cyclin-dependent protein kinase 5 (Cdk5) and the regulation of neurofilament metabolism. Eur J Biochem 268, 1534–1546 (2001).

40. M. Harterink et al., TRIM46 Organizes Microtubule Fasciculation in the Axon Initial Segment. J Neurosci 39, 4864–4873 (2019).

41. T. B. Shea et al., Cdk5 regulates axonal transport and phosphorylation of neurofilaments in cultured neurons. J Cell Sci 117, 933–941 (2004).

42. S. F. B. van Beuningen et al., TRIM46 Controls Neuronal Polarity and Axon Specification by Driving the Formation of Parallel Microtubule Arrays. Neuron 88, 1208–1226 (2015).

43. J. M. Brittain, Y. Wang, O. Eruvwetere, R. Khanna, Cdk5-mediated phosphorylation of CRMP-2 enhances its interaction with CaV2.2. FEBS Lett 586, 3813–3818 (2012).

44. R. Dong, X. Li, K. O. Lai, Activity and Function of the PRMT8 Protein Arginine Methyltransferase in Neurons. Life (Basel) 11 (2021).

45. T. Araki, J. Milbrandt, Ninjurin, a novel adhesion molecule, is induced by nerve injury and promotes axonal growth. Neuron 17, 353–361 (1996).

46. S. Marcos et al., Secreted frizzled related proteins modulate pathfinding and fasciculation of mouse retina ganglion cell axons by direct and indirect mechanisms. J Neurosci 35, 4729–4740 (2015).

47. J. Rodriguez et al., SFRP1 regulates the growth of retinal ganglion cell axons through the Fz2 receptor. Nat Neurosci 8, 1301–1309 (2005).

48. P. J. Hooikaas et al., MAP7 family proteins regulate kinesin-1 recruitment and activation. J Cell Biol 218, 1298–1318 (2019).

49. R. Kingston, D. Amin, S. Misra, J. M. Gross, T. Kuwajima, Serotonin transporter-mediated molecular axis regulates regional retinal ganglion cell vulnerability and axon regeneration after nerve injury. PLoS Genet 17, e1009885 (2021).

50. R. E. Hausman, Ocular extracellular matrices in development. Prog Retin Eye Res 26, 162–188 (2007).

51. S. F. Oster, D. W. Sretavan, Connecting the eye to the brain: the molecular basis of ganglion cell axon guidance. Br J Ophthalmol 87, 639–645 (2003).

52. C. Bonnans, J. Chou, Z. Werb, Remodelling the extracellular matrix in development and disease. Nat Rev Mol Cell Biol 15, 786–801 (2014).

53. J. Reinhard, S. Wiemann, S. Hildebrandt, A. Faissner, Extracellular Matrix Remodeling in the Retina and Optic Nerve of a Novel Glaucoma Mouse Model. Biology (Basel) 10 (2021).

54. D. M. Kim, C. M. Nimigean, Voltage-Gated Potassium Channels: A Structural Examination of Selectivity and Gating. Cold Spring Harb Perspect Biol 8 (2016).

55. N. S. Pollock, S. C. Ferguson, S. McFarlane, Expression of voltage-dependent potassium channels in the developing visual system of Xenopus laevis. J Comp Neurol 452, 381–391 (2002).

56. S. C. Ferguson, S. McFarlane, GABA and development of the Xenopus optic projection. J Neurobiol 51, 272–284 (2002).

57. J. W. Bang et al., GABA decrease is associated with degraded neural specificity in the visual cortex of glaucoma patients. Commun Biol 6, 679 (2023).

58. C. E. Mac Nair, R. W. Nickells, Neuroinflammation in Glaucoma and Optic Nerve Damage. Prog Mol Biol Transl Sci 134, 343–363 (2015).

59. C. Gomes et al., Induction of astrocyte reactivity promotes neurodegeneration in human pluripotent stem cell models. Stem Cell Reports 19, 1122–1136 (2024).

60. S. A. Liddelow et al., Neurotoxic reactive astrocytes are induced by activated microglia. Nature 541, 481–487 (2017).

61. L. Barbar et al., CD49f Is a Novel Marker of Functional and Reactive Human iPSC-Derived Astrocytes. Neuron 107, 436–453 e412 (2020).

62. P. Maddineni et al., CNS axonal degeneration and transport deficits at the optic nerve head precede structural and functional loss of retinal ganglion cells in a mouse model of glaucoma. Mol Neurodegener 15, 48 (2020).

63. L. Coughlin, R. S. Morrison, P. J. Horner, D. M. Inman, Mitochondrial morphology differences and mitophagy deficit in murine glaucomatous optic nerve. Invest Ophthalmol Vis Sci 56, 1437–1446 (2015).

64. Y. Liang et al., Axonal mitophagy in retinal ganglion cells. Cell Commun Signal 22, 382 (2024).

65. M. T. Maloney et al., Failure to Thrive: Impaired BDNF Transport along the Cortical-Striatal Axis in Mouse Q140 Neurons of Huntington’s Disease. Biology (Basel) 12 (2023).

66. H. A. Quigley et al., Retrograde axonal transport of BDNF in retinal ganglion cells is blocked by acute IOP elevation in rats. Invest Ophthalmol Vis Sci 41, 3460–3466 (2000).

67. O. Errea, B. Moreno, A. Gonzalez-Franquesa, P. M. Garcia-Roves, P. Villoslada, The disruption of mitochondrial axonal transport is an early event in neuroinflammation. J Neuroinflammation 12, 152 (2015).

68. C. M. Dengler-Crish et al., Anterograde transport blockade precedes deficits in retrograde transport in the visual projection of the DBA/2J mouse model of glaucoma. Front Neurosci 8, 290 (2014).

69. J. H. Liang et al., Dopamine signaling from ganglion cells directs layer-specific angiogenesis in the retina. Curr Biol 33, 3821–3834 e3825 (2023).

70. R. A. Warwick, A. S. Heukamp, S. Riccitelli, M. Rivlin-Etzion, Dopamine differentially affects retinal circuits to shape the retinal code. J Physiol 601, 1265–1286 (2023).

71. S. Yadav, P. J. Verma, D. Panda, C-terminal region of MAP7 domain containing protein 3 (MAP7D3) promotes microtubule polymerization by binding at the C-terminal tail of tubulin. PLoS One 9, e99539 (2014).

72. H. A. Quigley, E. M. Addicks, Regional differences in the structure of the lamina cribrosa and their relation to glaucomatous optic nerve damage. Arch Ophthalmol 99, 137–143 (1981).

73. I. A. Sigal, C. R. Ethier, Biomechanics of the optic nerve head. Exp Eye Res 88, 799–807 (2009).

74. K. Ono et al., Origin of Oligodendrocytes in the Vertebrate Optic Nerve: A Review. Neurochem Res 43, 3–11 (2018).

75. S. Hammer et al., Nuclei-specific differences in nerve terminal distribution, morphology, and development in mouse visual thalamus. Neural Dev 9, 16 (2014).

76. C. M. Fligor, K. C. Huang, S. S. Lavekar, K. B. VanderWall, J. S. Meyer, Differentiation of retinal organoids from human pluripotent stem cells. Methods Cell Biol 159, 279–302 (2020).

77. S. K. Ohlemacher, C. L. Iglesias, A. Sridhar, D. M. Gamm, J. S. Meyer, Generation of highly enriched populations of optic vesicle-like retinal cells from human pluripotent stem cells. Curr Protoc Stem Cell Biol 32, 1H 8 1–1H 8 20 (2015).

78. J. S. Meyer et al., Modeling early retinal development with human embryonic and induced pluripotent stem cells. Proc Natl Acad Sci U S A 106, 16698–16703 (2009).

79. R. Krencik, S. C. Zhang, Directed differentiation of functional astroglial subtypes from human pluripotent stem cells. Nat Protoc 6, 1710–1717 (2011).

80. C. M. Fligor et al., Extension of retinofugal projections in an assembled model of human pluripotent stem cell-derived organoids. Stem Cell Reports 16, 2228–2241 (2021).

81. A. Sridhar, M. M. Steward, J. S. Meyer, Nonxenogeneic growth and retinal differentiation of human induced pluripotent stem cells. Stem Cells Transl Med 2, 255–264 (2013).

82. Y. Mou, S. Mukte, E. Chai, J. Dein, X. J. Li, Analyzing Mitochondrial Transport and Morphology in Human Induced Pluripotent Stem Cell-Derived Neurons in Hereditary Spastic Paraplegia. J Vis Exp 10.3791/60548 (2020).

