## Supplemental Methods for "Modeling human retinal ganglion cell axonal outgrowth, development, and pathology using pluripotent stem cell-based microfluidic platforms"

**Maintenance and expansion of hPSCs**

hPSC lines used in this study included H7-BRN3b:tdTomato:Thy1.2 (1, 2) and H7-BRN3b:tdTomato:Thy1.2-OPTN(E50K), as described previously (1, 3). hPSC colonies were maintained on matrigel (Fisher Scientific, cat. #8774552) or Geltrex (Life Technologies, cat. #A1413302) coated plates in mTeSR1 (StemCell Technologies, cat. #85850) medium, with fresh media exchanged every day. At approximately 80% confluency, hPSC colonies were enzymatically lifted using dispase (2 mg/ml, Life Technologies, cat. #17105041), and expanded at a ratio of 1:6 or differentiation was initiated, as previous described (3).

**Retinal organoid differentiation and RGCs isolation**

hPSCs were differentiated into RGCs using previously published protocols (3-6). Briefly, embryoid bodies (EBs) were generated by lifting undifferentiated hPSC colonies with dispase (2 mg/ml), and resulting cell aggregates were slowly transitioned from mTeSR1 to neural induction media [NIM, DMEM/F12 with 1% N2 supplement, MEM nonessential amino acids, and heparin (2 ug/ml, Millipore-Sima, H3149)] over a period of 3 days. At day 6, EBs were fed with fresh NIM media and supplemented with bone morphogenetic protein 4 (BMP4, 50ng/mL, R&D Systems, 314-BP). At day 8, EBs were induced to adhere to a 6 well plate in the presence of 10% fetal bovine serum (FBS). Half-media changes with NIM were performed at day 9 and day 12, with a full media change at day 15. On day 16, presumptive retinal organoids were mechanically lifted and placed in suspension culture in Retinal Differentiation Medium (RDM, 1:1 DMEM:DMEM/F12, 2% B27 and 1% MEM NEAA), with half-media changes every other day. From day 20-35, the medium was gradually supplemented with FBS from 1-10% to enhance the survival of retinal organoids. On day 35, the medium was replaced by retinal maturation medium (RDM supplemented with 10% FBS, 1× GlutaMAX, and 100 μM Taurine), and half medium was changed every 2-3 days until day 45. At 45 days, retinal organoids were enzymatically dissociated with AccuMax (Life Technologies, cat. #00-4666-56) for 20 minutes at 37^o^C and RGCs were purified by Magnetic Activated Cell Sorting (MACS) using CD90 (Thy1.2) MicroBeads (Miltenyi Biotec, cat. #130-121-278), as previously described (1) and following the manufacturer’s protocol. Purified RGCs were plated on either poly-D-ornithine and laminin-coated glass coverslips (density of 450 cells/mm^2^) or pre-coated microfluidic devices and maintained in neuron maturation medium (1:1 of DMEM/F12 (1:1) and Neurobasal medium (Life Technologies, cat. #21103049) supplemented with 1× B27, 1× N2, 20 ng/mL BDNF, 20 ng/mL GDNF, 1× GlutaMAX and penicillin/streptomycin. Half-media was changed every 3-4 days for a total of 4 weeks for RGC maturation.

**Astrocyte differentiation and induction of a reactive state**

hPSCs were differentiated to an astrocyte lineage according to previously established protocols (7-9). Briefly, EBs were generated by lifting undifferentiated hPSC colonies with dispase, and resulting cell aggregates were slowly transitioned from mTeSR1 to neural induction media (NIM) over a period of 3 days. At day 7, EBs were plated on a laminin-coated 6-well plate with fresh NIM and fed every other day. At day 16, aggregates were mechanically lifted to form neurospheres and placed in differentiation medium (1:1 DMEM:DMEM/F12, 2% B27 and 1% MEM NEAA) in suspension culture. At day 40, neurospheres were supplemented with 20 ng/mL fibroblast growth factor 2 (FGF2, R&D Systems, 3718-FB) and 20 ng/mL epidermal growth factor (EGF, R&D Systems, 236-GMP). Neurospheres were mechanically passaged with a tissue chopper every 2 weeks for at least 6 months to bypass the neurogenic phase of differentiation and reach a gliogenic phase. To isolate astrocytes for experimental use, neurospheres were dissociated with AccuMax for 20 minutes at 37^o^C. Single cells were then plated on microfluidic devices in serum-free BrainPhys culture medium (StemCell Technologies, 5790) containing 2% B27, 1% N2, ascorbic acid, cAMP, BDNF (20ng/ml, Peprotech, 450-02), GDNF (20ng/ml, Peprotech, 450-10) and CNTF (20ng/ml, Peprotech, 450-13) for further maturation (8, 9). To induce a reactive phenotype, hPSC-derived astrocytes were incubated with TNFα (30 ng/mL, Peprotech), IL-1α (3 ng/mL, Peprotech), and C1q (400 ng/mL, MyBiosource), as previously described (10, 11).

**Cell seeding and maintenance of microfluidic devices**

Microfluidic devices were coated and handled following manufacturer’s instructions and as previously described (12). 2-compartment XonaChips (XC450) and 3-compartment XonaChips (XC-T500) from Xona microfluidics^®^ (North Carolina, USA) were used in this study. Briefly, the chips were coated with XC Pre-Coat^TM^ solution and washed twice with PBS, according to manufacturer’s instructions. The microfluidic chips were then coated with poly-D-ornithine (100µg/ml) overnight at 4°C. On the next day, the chips were washed twice with Milli-Q water and coated overnight with laminin at 37°C, 5% CO_2_. The cell seeding in 2- or 3-compartment microfluidics, as well as the differential volumes in the different chambers were as follows:

*2-compartment XonaChips (XC450)*: RGCs were plated within microfluidic devices at a density of 50,000 cells/devices (for imaging) or 250,000 cells/device (for RNA extraction) in the proximal chamber in soma media (retinal maturation medium). To induce axonal recruitment into the contralateral chamber of each device, extra BDNF (50 ng/ml) and CNTF (20 ng/ml) was added to all axonal chambers to recruit axons crossing over. Additionally, a differential volume between chambers was established, with the first chamber (soma) containing 200 µl of retinal maturation medium (described above), and the distal chamber (axon) containing 150 µl. hPSC-derived RGCs were maintained for 4 weeks, with media changes occurring every 2-3 days.

*3-compartment (XC-T500)*: RGCs were plated at a density of 150,000 cells/device into the proximal chamber. RGCs were grown in soma media for approximately 1.5 weeks before the seeding of astrocytes. hPSC-derived astrocytes were added to the middle chamber of each 3-compartment microfluidic at a density of 300,000 astrocytes (1:2 proportion of RGCs: astrocytes respectively) in axon media. In 3-compartment chamber, a differential volume between chambers was also established to recruit axons crossing over, with the proximal chamber (RGC somas) containing 250 µl of serum-free BrainPhys culture medium (StemCell Technologies, 5790) containing 2% B27, 1% N2, BDNF (20ng/ml, Peprotech, 450-02), GDNF (20ng/ml, Peprotech, 450-10) and CNTF (20ng/ml, Peprotech, 45013) (9), the middle chamber (astrocytes and RGC axons) contained 180 µl of axon media [BrainPhys culture medium supplemented with extra BDNF (50 ng/ml) and CNDF (20 ng/ml)], and the distal chamber (RGC axons) contained 150 µl of axon media. Cultures were maintained for maturation and axon extension for 2 weeks before the induction of astrocyte reactivity. To induce a reactive phenotype in hPSC-derived astrocytes, axonal media containing tumor necrosis factor alpha (TNFα, 30 ng/mL), interleukin-1 alpha (IL-1α, 3 ng/mL), and complement component 1q (C1q, 400 ng/mL) was added to the middle chamber every 2-3 days for 2 weeks (10). Cells were maintained in microfluid device for up to 5 weeks, with media changes every 2-3 days.

**Immunocytochemistry**

hPSC-derived RGCs seeded on coverslips or microfluidic devices were fixed with 4% paraformaldehyde at the indicated timepoints, and immunostained as previously described (6, 9, 13). Cells were fixed in 4% paraformaldehyde at room temperature (RT) for 30 minutes. After fixation, samples were washed 3 times with 1x PBS for 5 minutes each. Cells were then permeabilized in 0.2% Triton X-100 for 10 minutes and blocked in 1% BSA and 10% donkey serum for 1 hour, at RT. Primary antibodies, anti-MAP2 (rabbit, Santa Cruz #sc-20172, 1:200) and anti-SMI312 (mouse, BioLegend #83704, 1:250) were diluted in 0.5% BSA and 5% donkey serum and incubated on samples overnight at 4°C. The next day, samples were washed 3 times with 1x PBS, secondary antibodies were diluted on 0.5% BSA and 5% donkey serum for 1 hour at RT. Samples were then washed 3 times with 1x PBS and coverslips were mounted on slides using Fluoromount-G mounting medium. For microfluidic devices, 1x PBS was added to the wells for imaging. Imaging was performed using a Nikon A1R Confocal Microscope with Z-stack and/or tile scanning.

For the microfluidics containing astrocytes in the middle chamber, sequential immunostaining was performed. Briefly, after permeabilization and blocking, samples were incubated overnight at 4^o^C with anti-MAP2 (rabbit, Santa Cruz #sc-20172, 1:200) and anti-SMI312 (mouse, BioLegend #83704, 1:250). The next day, after the incubation with the secondary antibodies, samples were incubated overnight at 4^o^C with primary antibody against GFAP (rabbit, D1F4Q - Cell Signaling #12389S, 1:250). After the second overnight incubation, samples were stained with secondary antibody, washed 3 times with 1x PBS and imaged using a Nikon A1R Confocal Microscope with Z-stack.

**Dendritic and axonal quantification**

For morphological analysis and measurements, fluorescent images were analyzed using the ImageJ (Fiji) software. MAP2 staining in the somatic chamber was used to determine the number of primary dendrites as well as the soma size. For analysis of the axonal region, SMI-312 fluorescent images were converted into 8-bit image files and a threshold was used to determine the area occupied by the axons. Moreover, for morphological analysis, Sholl analysis plugin was used to determine axonal extension and complexity.

Imaris Image Analysis Software was used to determine the co-localization of MAP2-positive dendrites and SMI-312 axons in standard and microfluidic cultures. Within Imaris, MAP2 and SMI-312 were identified and isolated as surfaces using machine learning, and the surface of MAP2 was masked. Separate gray values were found for MAP2 and SMI-312, which were then used as their respective thresholds in the colocalization (coloc) module. Working in the coloc module, the region of interest (ROI) was set as the masked MAP2 surface with a threshold of 1000, and Channels A and B were set as MAP2 and SMI-312, respectively, with their thresholds included. Once completed, a coloc channel was created to identify the areas in which MAP2 and SMI-312 overlapped one another. Results are represented as the percentage of MAP2 co-localized with SMI-312 staining.

**Mitochondria transportation recording**

Mitochondria length and movement were determined in hPSC-derived RGCs, after 4 weeks of maturation, as previously described (14) with modifications. First, the medium of hPSC-RGCs was removed and cells were incubated with 50nM MitoTracker Green (Life technologies, cat #M7514) in fresh warm medium for 30 min at 37°C, 5% CO_2_. hPSC-RGCs were then washed three times with warm 1x PBS and BrainPhys Imaging Optimized Medium (StemCell Technologies, cat. #05796) supplement with 1´ B27, 1´ N2, and penicillin/streptomycin was added. Cells were maintained at 37°C, 5% CO_2_ for at least 20 min to recover prior imaging. Time lapse Imaging was performed using 40X oil lens in Nikon A1R Confocal Microscope with 5 min scanning equipped with CO_2_ chamber. The generation of kymograph and movement quantification were performed in Fiji, as described (14).

**RNA isolation and mRNA sequencing**

At 4 weeks after plating, RNA was collected from soma and axon compartments of microfluidic devices using the PicoPure^TM^ RNA Isolation Kit (KIT0204, Applied Biosystems), following manufacturer’s instructions, and as previously described (15). Due to the low RNA concentration obtained from the axon compartment, samples were then pre-amplified using the SuperScript™ IV Single Cell/Low Input cDNA PreAmp Kit (Invitrogen, 11752048), following manufacturer’s instructions. After pre-amplification, cDNA samples were purified using the Agencourt AMPure XP beads (Beckman Coulter, A63880) and the DynaMag-2 Magnet (Invitrogen, 12321D). The cDNA products were next checked on Agilent Bioanalyzer. The Nextera XT DNA Library Preparation Kit (Illumina) was used for subsequent library preparation. The resulting libraries were quantified, and quality accessed by Qubit and Agilent TapeStation. The libraries were pooled in equal molarity and sequenced with 2×150bp paired-end configuration on an Illumina NovaSeq 6000 sequencer.

**RNA-sequencing analysis**

The sequencing data was first processed using FastQC (Babraham Bioinfomatics, Cambridge, UK) and then mapped to the Human genome (GRCH38) using STAR RNA-seq aligner (54) with the parameter: “— outSAMmapqUNIQUE 60”. Uniquely mapped sequencing reads were assigned to GRCH38 reference genome using featureCounts. Differentially expressed genes were tested by using DESeq2 with pvalue<0.05 as the significant cutoff (16) for further pathway analysis. Pathway enrichment analysis were conducted by hypergeometric test against Human Gene Ontology and MsigDB v6 canonical pathways, with pvalue<0.05 as the significant cutoff (17).

**Statistical analysis**

For all studies a minimum of at least three biological replicates were used in each assay. Differences between isogenic control and OPTN(E50K) RGCs, or between RGCs co-cultured with control or reactive astrocytes, were determined by two-tailed Student’s t-test analysis. ANOVA was used when more than two groups were considered. The statistical test used for each result is described in the legend of the respective figure. Statistical analyses were performed using GraphPad PRISM 9.5.0. p < 0.05 was considered statistically significant.

**REFERENCES**

1.	V. M. Sluch *et al.*, Enhanced Stem Cell Differentiation and Immunopurification of Genome Engineered Human Retinal Ganglion Cells. *Stem Cells Transl Med* **6**, 1972-1986 (2017).

2.	J. A. Thomson *et al.*, Embryonic stem cell lines derived from human blastocysts. *Science* **282**, 1145-1147 (1998).

3.	K. B. VanderWall *et al.*, Retinal Ganglion Cells With a Glaucoma OPTN(E50K) Mutation Exhibit Neurodegenerative Phenotypes when Derived from Three-Dimensional Retinal Organoids. *Stem Cell Reports* **15**, 52-66 (2020).

4.	C. M. Fligor, K. C. Huang, S. S. Lavekar, K. B. VanderWall, J. S. Meyer, Differentiation of retinal organoids from human pluripotent stem cells. *Methods Cell Biol* **159**, 279-302 (2020).

5.	S. K. Ohlemacher, C. L. Iglesias, A. Sridhar, D. M. Gamm, J. S. Meyer, Generation of highly enriched populations of optic vesicle-like retinal cells from human pluripotent stem cells. *Curr Protoc Stem Cell Biol* **32**, 1H 8 1-1H 8 20 (2015).

6.	J. S. Meyer *et al.*, Modeling early retinal development with human embryonic and induced pluripotent stem cells. *Proc Natl Acad Sci U S A* **106**, 16698-16703 (2009).

7.	R. Krencik, S. C. Zhang, Directed differentiation of functional astroglial subtypes from human pluripotent stem cells. *Nat Protoc* **6**, 1710-1717 (2011).

8.	K. B. VanderWall *et al.*, Astrocytes Regulate the Development and Maturation of Retinal Ganglion Cells Derived from Human Pluripotent Stem Cells. *Stem Cell Reports* **12**, 201-212 (2019).

9.	C. Gomes *et al.*, Astrocytes modulate neurodegenerative phenotypes associated with glaucoma in OPTN(E50K) human stem cell-derived retinal ganglion cells. *Stem Cell Reports* **17**, 1636-1649 (2022).

10.	S. A. Liddelow *et al.*, Neurotoxic reactive astrocytes are induced by activated microglia. *Nature* **541**, 481-487 (2017).

11.	L. Barbar *et al.*, CD49f Is a Novel Marker of Functional and Reactive Human iPSC-Derived Astrocytes. *Neuron* **107**, 436-453 e412 (2020).

12.	C. M. Fligor *et al.*, Extension of retinofugal projections in an assembled model of human pluripotent stem cell-derived organoids. *Stem Cell Reports* **16**, 2228-2241 (2021).

13.	A. Sridhar, M. M. Steward, J. S. Meyer, Nonxenogeneic growth and retinal differentiation of human induced pluripotent stem cells. *Stem Cells Transl Med* **2**, 255-264 (2013).

14.	Y. Mou, S. Mukte, E. Chai, J. Dein, X. J. Li, Analyzing Mitochondrial Transport and Morphology in Human Induced Pluripotent Stem Cell-Derived Neurons in Hereditary Spastic Paraplegia. *J Vis Exp* 10.3791/60548 (2020).

15.	J. Harkin *et al.*, A highly reproducible and efficient method for retinal organoid differentiation from human pluripotent stem cells. *Proc Natl Acad Sci U S A* **121**, e2317285121 (2024).

16.	M. I. Love, W. Huber, S. Anders, Moderated estimation of fold change and dispersion for RNA-seq data with DESeq2. *Genome Biol* **15**, 550 (2014).

17.	A. Subramanian *et al.*, Gene set enrichment analysis: a knowledge-based approach for interpreting genome-wide expression profiles. *Proc Natl Acad Sci U S A* **102**, 15545-15550 (2005).
